# Mild hyperthermia-induced thermogenesis in the endoplasmic reticulum defines stress response mechanisms

**DOI:** 10.1101/2024.05.24.595680

**Authors:** Barbara Dukic, Zsófia Ruppert, Melinda E. Tóth, Ákos Hunya, Ágnes Czibula, Péter Bíró, Ádám Tiszlavicz, Mária Péter, Gábor Balogh, Miklós Erdélyi, Gyula Timinszky, László Vígh, Imre Gombos, Zsolt Török

## Abstract

Previous studies reported that a mild, non-protein denaturing, fever-like temperature increase induced the unfolded protein response (UPR) in mammalian cells. Our dSTORM super-resolution microscopy experiments revealed that the master regulator of the UPR, the IRE1 (inositol-requiring enzyme 1) protein is clustered as a result of UPR activation in a human osteosarcoma cell line (U2OS) upon mild heat stress. Using ER thermo yellow, a temperature-sensitive fluorescent probe targeted to the endoplasmic reticulum (ER), we detected significant intracellular thermogenesis in mouse embryonic fibroblast (MEF) cells. Temperatures reached at least 8°C higher than the external environment (40°C), resulting in exceptionally high ER temperatures similar to those previously described for mitochondria. Mild heat-induced thermogenesis in the ER of MEF cells was likely due to the uncoupling of the Ca^2+^/ATPase (SERCA) pump. The high ER temperatures initiated a pronounced cytosolic heat shock response in MEF cells, which was significantly lower in U2OS cells in which both the ER thermogenesis and SERCA pump uncoupling were absent. Our results suggest that, depending on intrinsic cellular properties, mild hyperthemia-induced intracellular thermogenesis defines the cellular response mechanism, and determines the outcome of hyperthermic stress.

## Introduction

Eukaryotic cells have two evolutionarily highly conserved systems to cope with environmental or pathophysiological stress conditions: the heat shock response (HSR) and the unfolded protein response (UPR) (Almanza et al., 2019; Chen et al., 2023; Ron and Walter, 2007) The effects of stress depend on the stress dose such as intensity and duration. By characterizing its effect on Chinese Hamster Ovary (CHO) cells, we have previously classified heat stress into three distinct categories, namely: **severe**, involving induction of heat shock proteins (HSPs) together with major macromolecular damage or even cell death; **moderate** (negligible protein denaturation with less intense HSP induction), and **mild**, eliciting ‘eustress’ (without HSP induction) (Peksel et al., 2017). We also showed that mild heat triggers a distinct, dose-dependent remodeling of the cellular lipidome followed by the expression of HSPs only at higher temperatures (Tiszlavicz et al., 2022). A significant elevation in the relative concentration of saturated membrane lipid species and specific lysophosphatidylinositol and sphingolipid species suggests rapid membrane microdomain reorganization and an overall time-dependent increase in membrane rigidity in response to fluidizing heat. Our RNAseq experiments revealed that mild heat initiated stress-related signaling cascades in the endoplasmic reticulum (ER) in agreement with previous reports (Bettaieb and Averill-Bates, 2015; Tiszlavicz et al., 2022; Xu et al., 2011) resulting in dose-dependent lipid rearrangement and an elevated resistance to membrane fluidization by benzyl alcohol (Tiszlavicz et al., 2022). To protect cells against lethal, protein-denaturing high temperatures, the classical HSP response was evolved. The presence of distinct layers of stress response elicited by different heat dosages highlight the capability of cells to utilize multiple tools to resist and survive potentially lethal stress conditions.

The ER consists of a dynamic membrane network that is required for the synthesis and modification of proteins and lipids. The accumulation of unfolded proteins in the ER lumen activates an adaptive unfolded protein response, UPR, as a mechanism to restore homeostasis. The UPR is mediated by three ER-localized transmembrane proteins: inositol-requiring 1a (IRE1a), PKR-like endoplasmic reticulum kinase (PERK), and activating transcription factor 6 (ATF6) (Adams et al., 2019; Lindholm et al., 2017; Radanović and Ernst, 2021; Ron and Walter, 2007). Interestingly, recent findings show that lipid perturbation is also a direct activator of the UPR, independent of protein unfolding (Ernst et al., 2018; Fun and Thibault, 2020; Gianfrancesco et al., 2018; Halbleib et al., 2017; Tam et al., 2018; Xu and Taubert, 2021). Although the mechanism of the UPR process has been extensively investigated, the relationship between heat stress, especially mild hyperthermia, and the ER homeostatic response remains unclear.

Our previous observation that mild, non-protein-denaturing heat induced a type of UPR in mammalian cells (Tiszlavicz et al., 2022), initiated our studies on intracellular heat production. At the subcellular level, the mitochondrion is the organelle best-known for thermogenesis (Beignon et al., 2022). Using a temperature-sensitive fluorescent indicator targeted to mitochondria, it has been shown that mitochondria are physiologically maintained at close to 50°C (Chrétien et al., 2018). This high organelle temperature is maintained in the face of a variety of metabolic stresses, including substrate starvation or modification, or decreased ATP demand (Terzioglu et al., 2023). The ER has also gained attention as another kind of organelle responsible for heat production, mediated by a Ca^2+^-ATPase (SERCA) pump (Arai et al., 2014). Although these studies were not the first suggesting that mammalian mitochondria and the ER could be warmer than their surroundings, such discoveries fostered attention and discussion, especially considering the thermal physics of the phenomenon (Macherel et al., 2021). While there is still continuing debate on sizzling organelles, we aimed to investigate the intriguing possibility that the high metabolic demand of the cellular stress response could generate thermogenesis in the ER, further elevating its temperature and ultimately leading to a heat-shock response.

## Results

### Distinct cell types show differential stress transcriptome profiles upon heat treatment

The transcription profiles of key HSP and UPR genes were described in MEF and U2OS cells in response to heat treatment for 1 h at different temperatures (37°C, 39°C, 40°C, 41°C, 42°C, 43°C, 44°C) (**Fig. 1, Suppl. Table 1, Suppl. Fig. 1 and 2**). RT-qPCR analysis revealed a distinct, cell type-dependent response for the different levels of heat stress.

**Figure 1.**
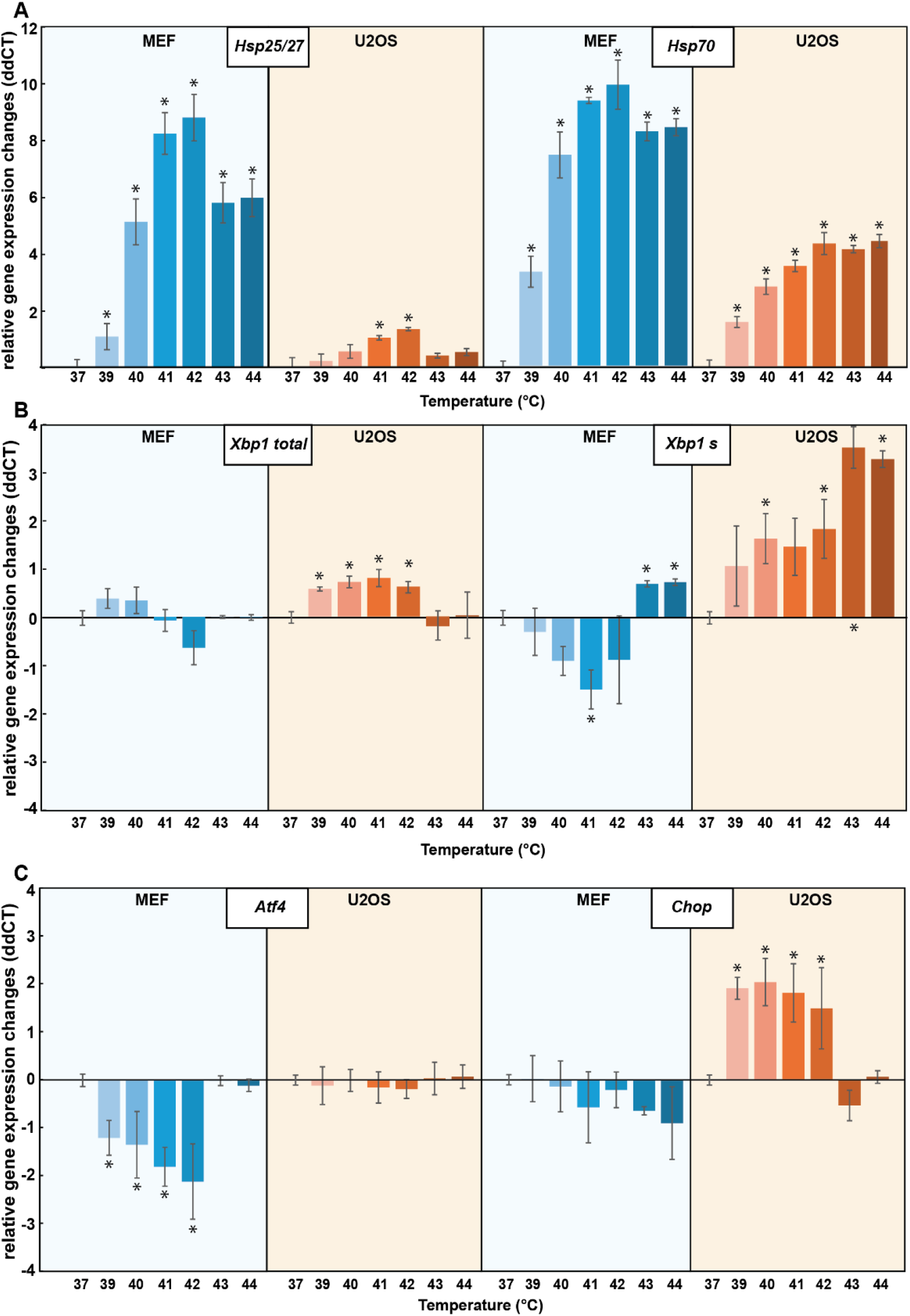
Relative gene expression changes (ddCT) in MEF and U2OS cells in response to heat treatment. The cells were subjected to different temperatures (39°C, 40°C, 41°C, 42°C, 43°C, 44°C) for 1 h or left untreated at 37°C. The relative expression of different (A) heat shock protein (HSP) and (B-C) unfolded protein response (UPR) genes was studied using RT-qPCR (*n* = 3 biological repetitions). Student’s t-test was used for statistical comparisons. Data represent mean ± SD, n = 3, \**p*<0.05 compared to 37°C.

In MEF cells, *Hsp25* and *Hsp70* genes exhibited significant upregulation already at mild (39-40°C) temperatures, while in U2OS cells the elevated expression of *HSP25* could only be observed at higher (41-42°C) temperatures (**Fig. 1A**). Transcription of the HSP genes was much higher in MEF cells than in U2OS cells, while the expression of the HSR master regulator HSF1 was significantly increased only in U2OS cells (**Suppl. Table 1 and Suppl. Fig. 1**).

To investigate the mechanism of the UPR, we focused on its three main signaling branches, represented by the *IRE1, PERK* and *ATF6* genes. Relative expression of these genes, along with *XBP1* (X-box binding protein 1), the downstream target of IRE1, and *ATF4* and *CHOP*, the targets of PERK, were analyzed. In U2OS cells, there was no change in the expression of *IRE1, PERK* and *ATF6* (**Suppl. Table 1 and Suppl. Fig. 2**), but our results revealed a significant increase in the expression of *XBP1* and *CHOP* upon heat treatment at 39-40°C (**Fig. 1B,C**). Notably, PERK-dependent *Atf4* expression was significantly downregulated in MEF cells at mild temperatures (39-42°C), and normalized under more severe heat conditions (43-44°C) (**Fig. 1C**).

Upon activation, IRE1 undergoes autophosphorylation and catalyzes RNase-mediated unconventional splicing of the XBP1 mRNA (Belyy et al., 2022). Our RT-qPCR results in U2OS cells revealed an increase in total *XBP1* expression, concomitant with an increased amount of spliced *XBP1* (*XBP1*-s) mRNA (**Suppl. Table 1 and Fig. 2B**). In parallel, the amount of unspliced *XBP1* (*XBP1*-us) slightly increased in the mild temperature region (39-41°C), but behaved oppositely in response to more severe heat treatment (43-44°C) by displaying a much more pronounced decrease (**Suppl. Fig. 2**). In MEF cells, there was no alteration either in the total or in the unspliced *Xbp1* level under mild heat stress (39-42°C), while at higher temperatures (43-44°C) we observed downregulation of the unspliced *Xbp1* and upregulation of the spliced form.

Pearson correlation analysis was conducted to examine the relationship between the induced HSR and UPR genes upon exposure of MEF and U2OS cells to heat, and distinct correlation patterns were observed (**Suppl. Table2, Suppl. Fig. 3**). MEF cells exhibited moderate to very strong positive correlations between the inductions of the examined HSP family members (r = 0.96 for *Hsp25* vs *Hsp70*). In U2OS cells, while the less pronounced *HSP* gene expression was paralleled by weaker correlations (r = 0.65 for *HSP25* vs *HSP70*). In contrast, the association was strong between certain UPR genes in U2OS but not in MEF cells: for *Xbp1 total* vs *Chop*, r = 0.84 in U2OS and r = 0.32 in MEF. On the other hand, we found a strong negative correlation between the induction of HSR and UPR genes in MEF cells (r = -0.9 for *Hsp90aa1* vs *Atf4*), whereas the same association was moderately positive in U2OS cells (r = 0.4) (**Suppl. Table 2 and Suppl. Fig. 3**).

### IRE1 clustering in response to mild heat treatment of 40°C

IRE1, one of the key UPR signal activators, is known to assemble into larger oligomers in response to treatment with tunicamycin, which induces ER stress (Belyy et al., 2020). In view of this finding, we asked whether a similar IRE1 clustering would result from mild, fever-like temperature stress. U2OS IRE1-mEOS cells, in which IRE1 transmembrane proteins were tagged with a fixation-resistant photoactivatable fluorescent mEos4 tag, were treated at 40°C or 42°C for one hour, and fixed. Imaging by dSTORM super-resolution microscopy revealed that the number of small clusters (< 5000 nm^2^) decreased, while the number of larger clusters increased in response to the heat treatments (**Fig. 2**). The increase in cluster area was highly significant at 40°C, and may be correlated with enhanced UPR signaling activity.

**Figure 2.**
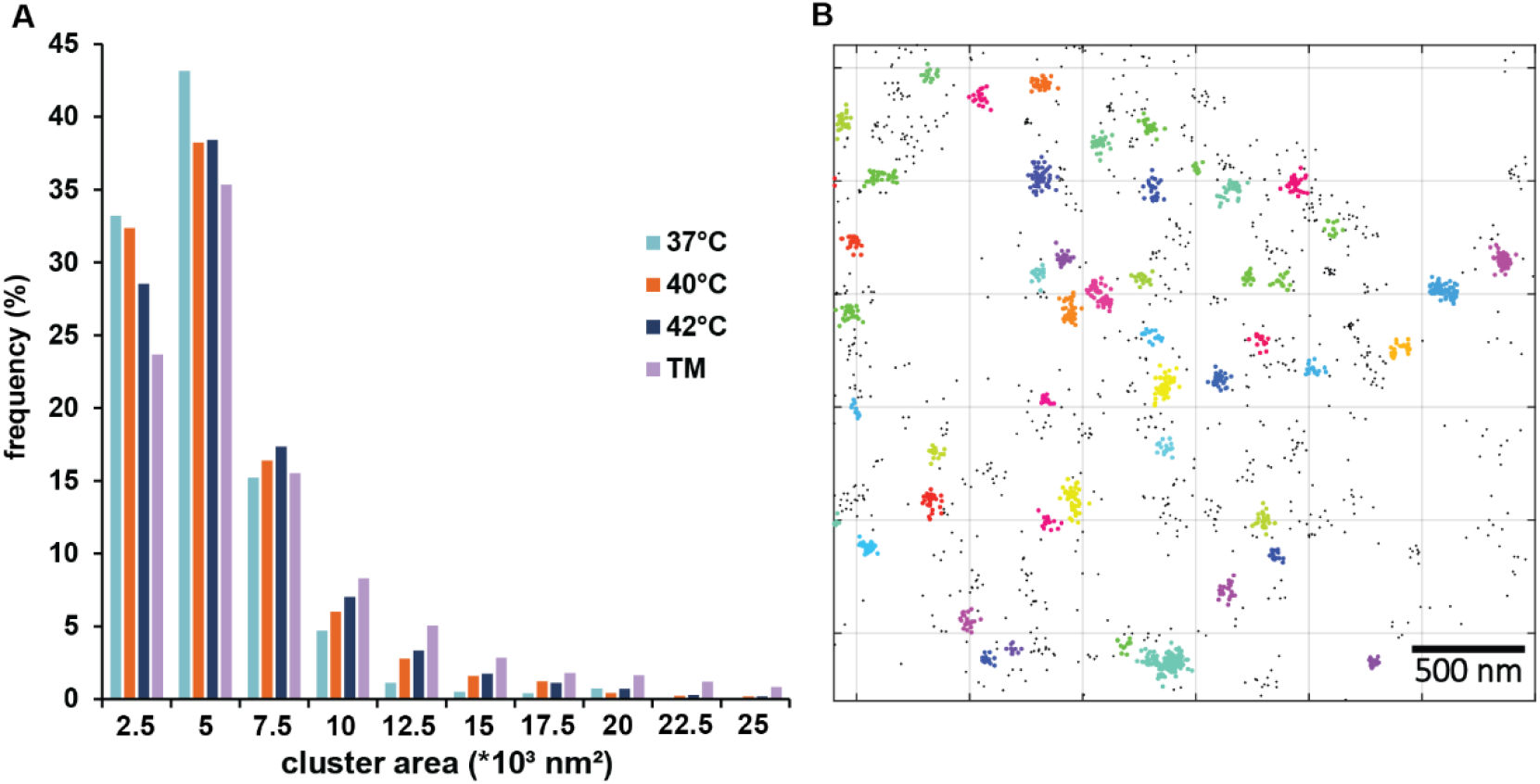
Clustering of the IRE1 transmembrane protein in fixed U2OS IRE1-mEOS cells upon heat treatment (1 h at 40°C or 42°C). (A) Size distribution of IRE1 clusters in non-treated (37°C) and heat-treated (40° or 42°C) U2OS cells. The Kolmogorov–Smirnov test was performed to analyze the equality of distributions. Samples treated at 40° or 42°C differed from the control (37°C) significantly. Tunicamycin (TM) treatment (5 μg/mL, 1 h) was used as the positive control. (B) Clusters of IRE1 molecules are labelled with colors in a representative dSTORM image after DBSCAN analysis (**see Suppl. Fig. 4**).

### IRE1-dependent XBP1 expression is upregulated upon mild heat treatment

To further investigate the intricate involvement of IRE1 in UPR signaling, IRE1-dependent XBP1 protein levels were measured during heat treatments using a fluorescence reporter construct in U2OS cells (Nougarède et al., 2018). Flow cytometry demonstrated significant increases in IRE1-dependent XBP1-mNeonGreen levels in cells subjected to mild heat treatments (40°C and 42°C, **Fig. 3**). These observations supported the results of RT-qPCR and dSTORM analyses and corroborated the activation of the UPR pathway under fever-like conditions.

**Figure 3.**
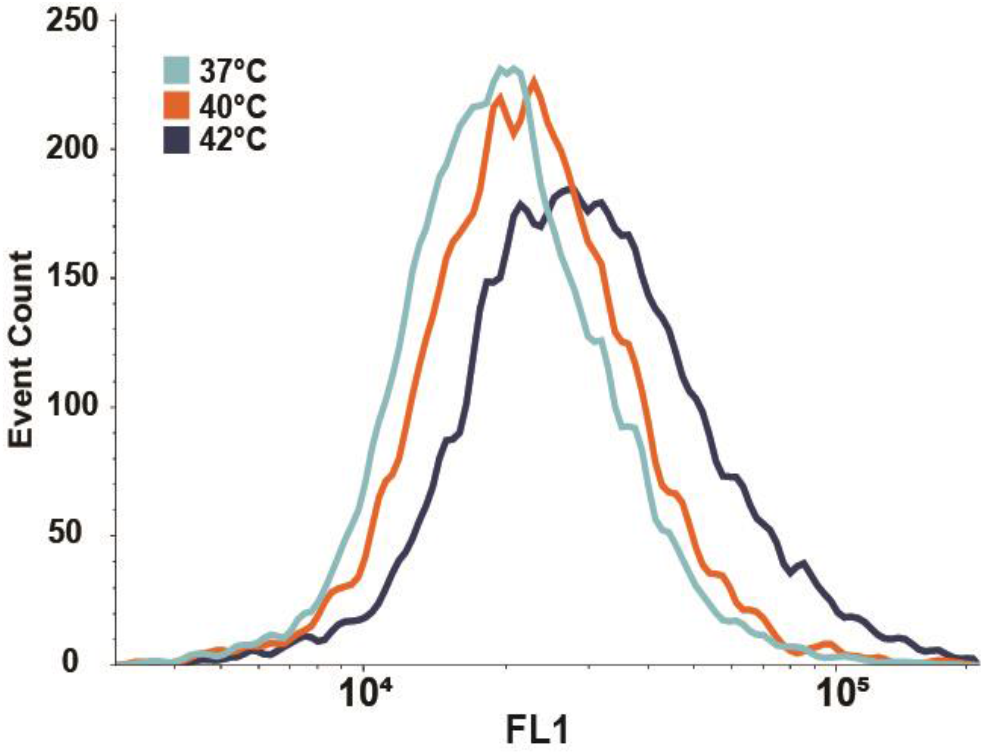
Flow cytometric determination of the level of XBP1 (mNeonGreen) protein in heat-stressed U2OS cells. Cells were heat stressed at 40°C or 42°C for one hour followed by six hours of recovery at 37°C. The Kolmogorov–Smirnov test was performed to analyze the equality of distributions. Samples treated at 40 or 42°C differed significantly from the control (37°C).

### Mild heat could induce thermogenesis in the ER

According to our previous findings and a literature search, the induction of HSP genes is a response typically associated with higher temperatures. To explain the unexpected increase in the induction of HSP genes in MEF cells elicited by mild heat, we proposed intracellular thermogenesis as the potential source. To test this hypothesis, we used ER thermo yellow, a fluorescent probe selectively targeting the ER, to measure the intracellular temperature. Since the probe’s intensity is not affected by enzymatic activity, metabolic activity, and cytosolic Ca^2+^ concentration, a decrease in the probe’s fluorescence intensity indicates increased temperature in the ER (Arai et al., 2014).

High-resolution fluorescence microscopy images of the ER thermo yellow-labeled MEF and U2OS cells (heat-stressed while on the microscope stage) showed significant differences in ER temperature of heat-treated cells (**Fig. 4A**). In live U2OS cells, the 40°C treatment did not increase the ER temperature more than in fixed cells. In contrast, in MEF cells a significant ER temperature elevation was observed. A calibration curve was used to estimate the extent of temperature increase, calculated from intensity changes of the fixed cells (**Fig. 4C**). This estimation revealed temperatures reaching at least 8°C higher than the external environment (40 °C) in the ER of live MEF cells.

**Figure 4.**
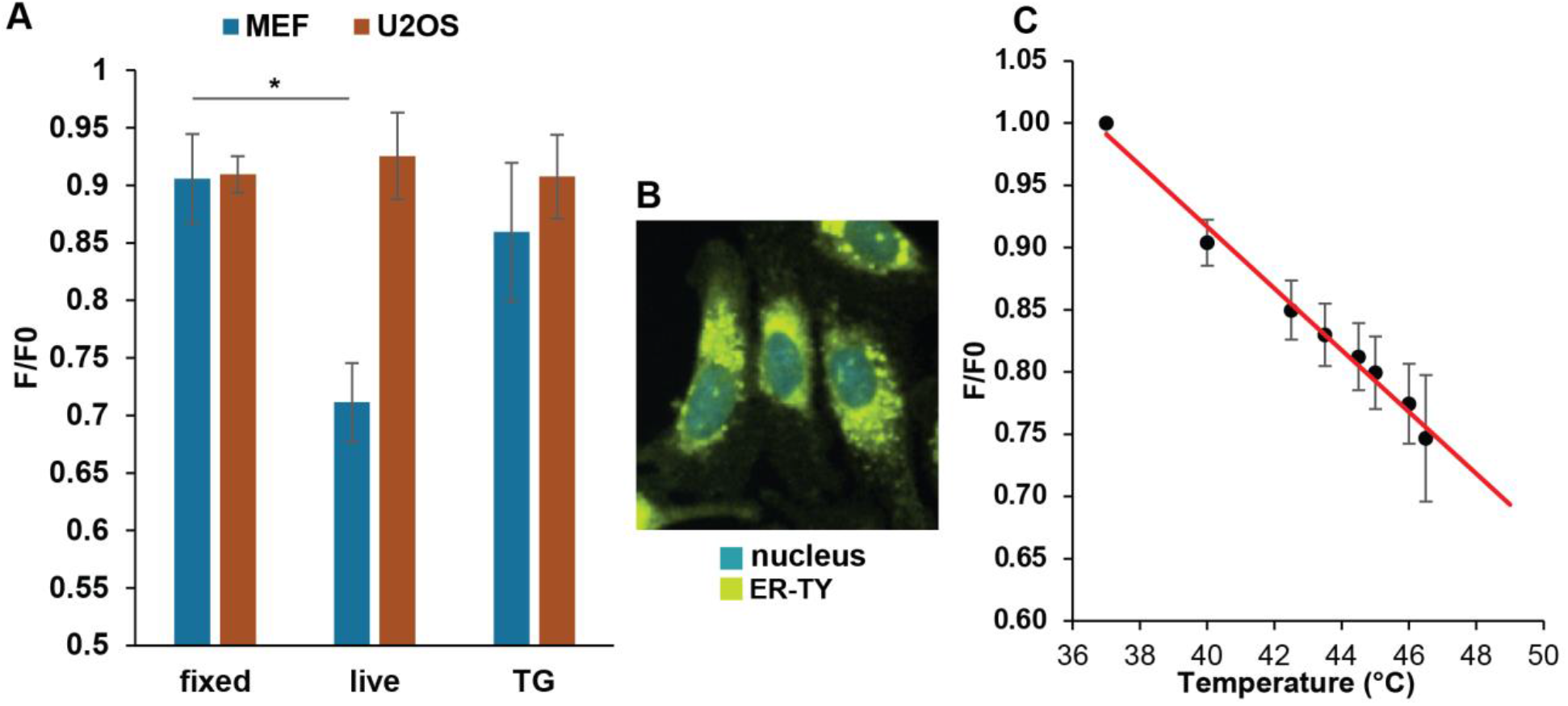
Effect of mild hyperthermia on the endoplasmic reticulum temperature in MEF and U2OS cells. (A) Changes in F/F0 ratio in live, fixed and 1 μM thapsigargin-pretreated cells upon 40°C heat treatment. The average fluorescence intensity values of the heat-treated sample, F were normalized with the average intensity values of the sample at 37°C, before the heat treatment, F0, by calculating the F/F0 ratio. For F/F0 = 1, there was no change in ER temperature. F/F0< 1 indicated increased ER temperature and F/F0 > 1 indicated decreased ER temperature. Student’s t-test was used for statistical comparisons. Data are mean ± SD, n = 1000, \**P*<0.05 compared to fixed cells. (B) Overlay of fluorescence images (blue, Hoechst 33342; yellow, ER thermo yellow). (C) Calibration curve calculated from the intensity changes of fixed cells (n = 10 cells).

The elevated ER temperature in MEF cells might result from SERCA pump uncoupling. In this state, the pump continues hydrolyzing ATP, but without effectively transporting calcium ions across the ER membrane, with the remaining energy dissipating as heat. To test this possibility, we treated groups of cells with the non-competitive SERCA inhibitor, thapsigargin, as a positive control (Meis, 2002). Thapsigargin treatment eliminated the increase in ER temperature in the MEF cells (**Fig. 4A**) suggesting that SERCA activity was indeed the source of the mild heat-induced ER thermogenesis.

### ER Ca^2+^ levels decrease upon mild heat treatment

To further support the mild heat-induced uncoupling of the SERCA pump in MEF but not in U2OS cells, we measured ER Ca^2+^ levels in both cell lines after 40°C heat treatment. The low affinity calcium probe, Mag-fluo-4, registers changes in Ca^2+^ level in a cell by changes in fluorescent intensity as measured by flow cytometry. Upon heat treatment, a significant decrease in ER Ca^2+^ levels was detected in MEF cells, contrasted with the absence of detectable changes in Ca^2+^ levels in U2OS cells (**Fig. 5**). This observation further strengthens our hypothesis that the observed temperature change in MEF cells is linked to SERCA pump uncoupling caused by fever-like heat treatment.

**Figure 5.**
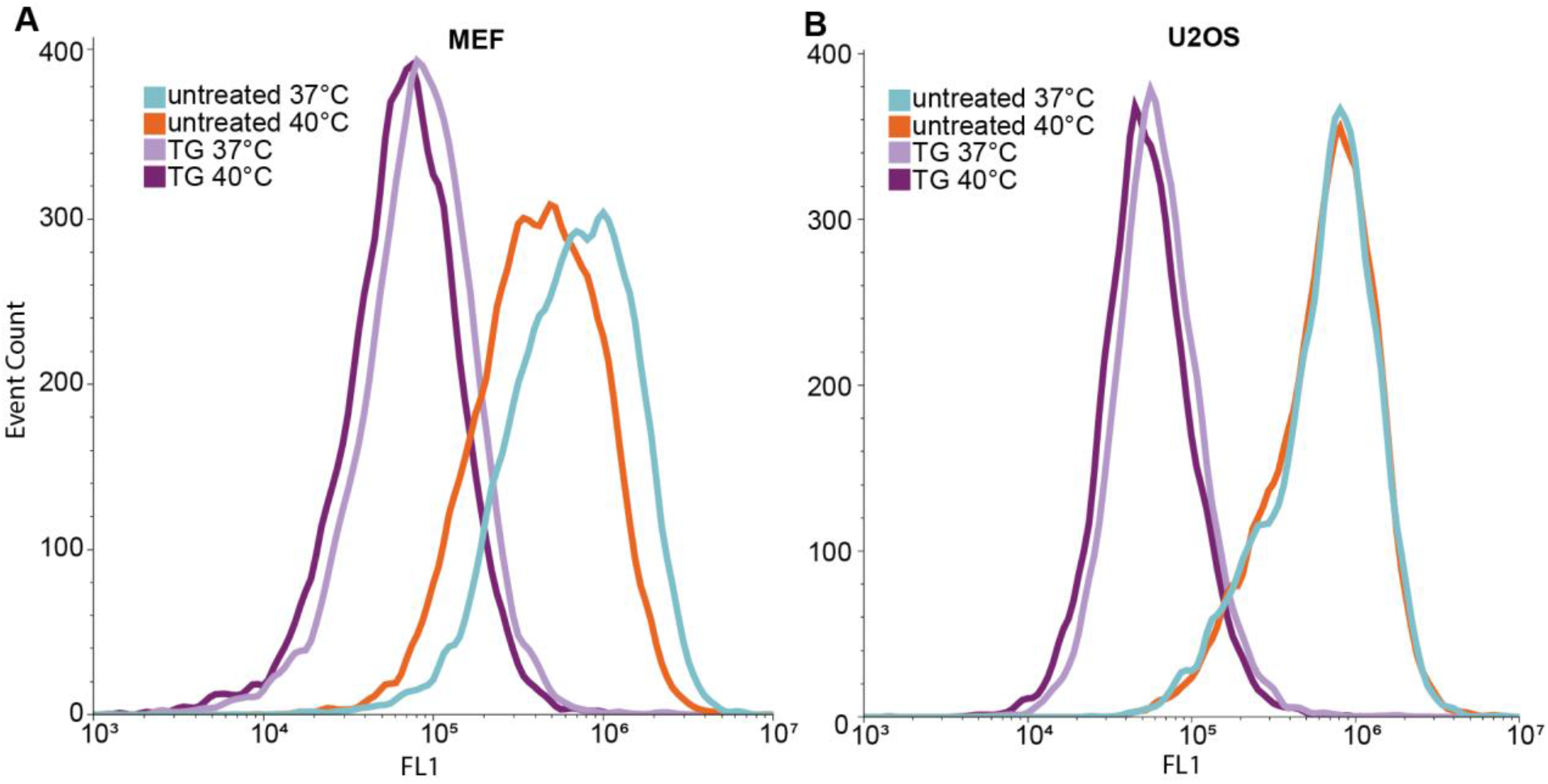
Effect of mild heat on endoplasmic reticulum Ca^2+^ levels. Flow cytometric analysis of endoplasmic reticulum Ca^2+^ levels in Mag-fluo-4-labeled, non-treated (37°C), heat-treated (40°C), and TG-pretreated (1 μM, 1 h) MEF and U2OS cells. The Kolmogorov-Smirnov test was performed to analyze the equality of distributions. MEF samples treated at 40° or 42°C differed from the control (37°C) significantly. TG: thapsigargin

Cells were also pre-treated with thapsigargin, which, as expected, resulted in decreased ER Ca^2+^ levels, disrupting cellular calcium homeostasis. TG-treated cells were then subjected to mild heat stress. Inhibition of the SERCA pump by thapsigargin prevented further Ca^2+^ depletion by heat, thus supporting the role of SERCA in intracellular heat production upon mild heat stress.

## Discussion

### Interplay between the UPR and HSR upon heat stress

Previously we described how Chinese hamster ovary (CHO) cells use distinct layers of stress response to protect against different levels of environmental stress (Tiszlavicz et al., 2022). We also identified the UPR as a first line of defense during mild hyperthermia resulting in membrane rearrangement and protection against various environmental challenges (Peksel et al., 2017; Tiszlavicz et al., 2022). To further generalize our previous findings, we used real-time quantitative PCR and super-resolution fluorescence microscopy to dissect the interrelationship and the mechanism of the switch between UPR and HSR in mouse embryonic fibroblast (MEF) and human osteosarcoma (U2OS) cell lines. By applying a range of heating challenges from mild to more severe (39-42°C) we detected the induction of both types of cellular defense, UPR and HSR, but with distinctive, cell-specific mechanisms. While UPR seems to be sufficient for withstanding mild hyperthermia in U2OS cells, this is not the case for MEF cells, where the HSR is induced at much lower temperatures (**Fig. 1**). Among the three branches of the UPR, the *IRE1* (XBP1) and *PERK* (CHOP, ATF4) pathways were activated by mild heat, while expression of the *ATF6* branch seemed to be silent in U2OS cells (**Suppl. Table 1 and Suppl. Fig. 2**). Previously it was shown for AD293 cells that a longer (1-16 hours) heat stress at 40°C could induce the downstream targets of ATF6; however, they were not induced for other cell types tested including two hepatocyte cell lines, a pancreatic beta cell line, and MEF cells (Xu et al., 2011). As ATF6 activation is influenced by its oxidation state, glycosylation state and proteosome-dependent turnover, it seems that stimuli preferentially affecting these aspects of ER protein processing do not occur during mild, fever-like hyperthermia but only with higher temperatures (Rutkowski and Hegde, 2010).

Pearson analysis has shown a negative correlation between the expression (including splicing) of specific HSR and UPR genes in MEF cells, with particularly strong correlation between the levels of *Hsp90aa1* mRNA and UPR mRNAs of *Xbp1_s* (r = 0.8) and *Atf4* (r = -0.9) (**Suppl. Table 2/Suppl. Fig. 3**). Since HSP90 was shown to stabilize both PERK and IRE1 (but not ATF6) by associating with their cytosolic domains it is possible that this phenomenon is the result of the dissociation of HSP90 from the two ER sensor kinases stimulated by intensive intracellular thermogenesis (Marcu et al., 2002; Somogyvári et al., 2022). It was reported earlier for five different mammalian cell types that exposure of cells to longer periods of hyperthermia at 40°C resulted in a partial or full ER stress pathway induction as well as a HSR, while at 43°C the ER stress pathway was inhibited (Xu et al., 2011). In contrast, our results indicate that under extreme conditions when the stress cannot be overcome, a different pathway of the UPR can be activated for the purpose of initiating apoptosis, cell death and clearance (**Fig. 6**) (Walczak et al., 2019). These observations suggest that there are first and second waves in the UPR, depending on the severity of the stress (**Fig. 6**).

**Figure 6.**
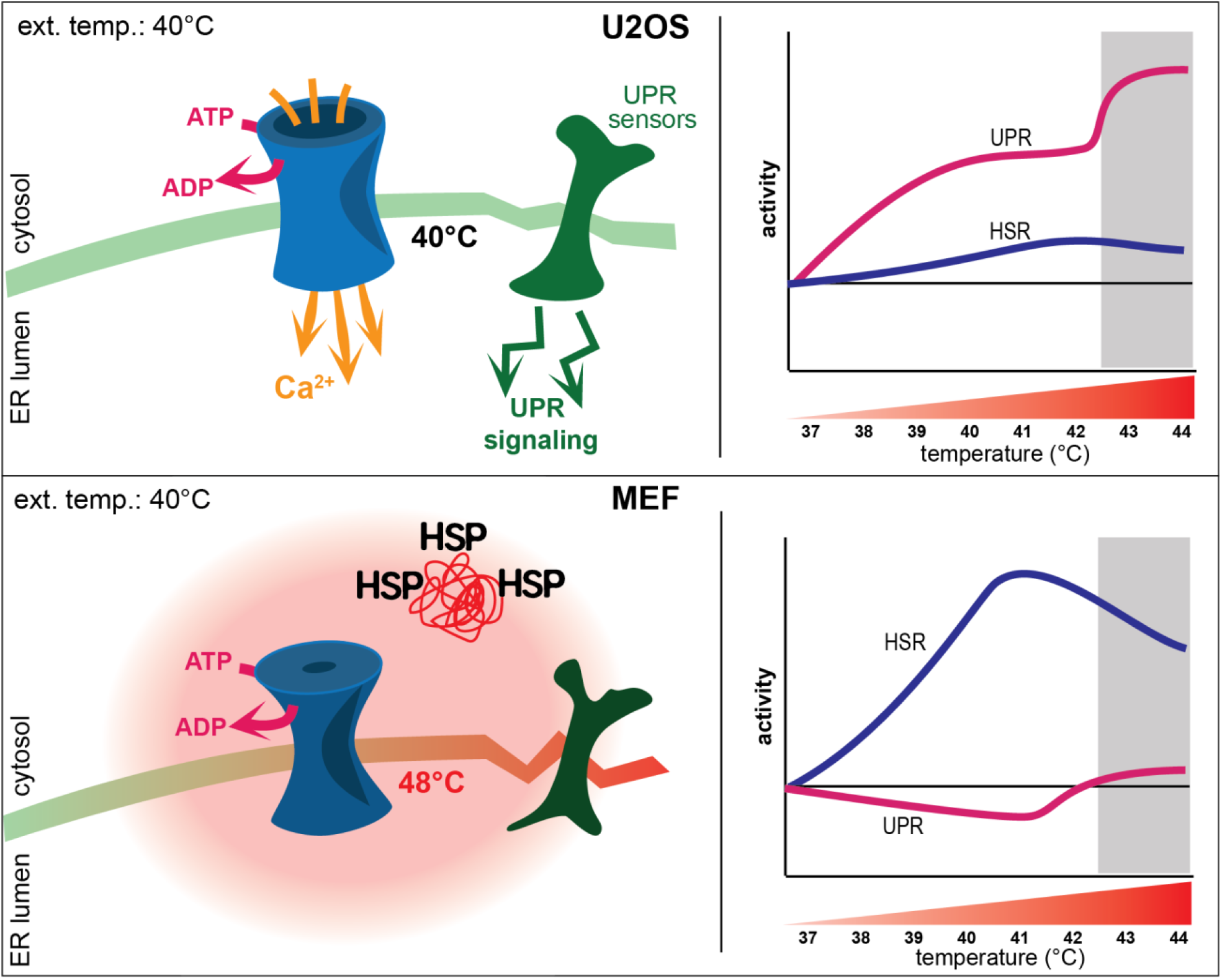
Cell-specific alternative stress response mechanisms. Under mild hyperthermia, U2OS cells implement both the UPR and HSR to maintain membrane and protein homeostasis, while under more severe heat conditions (above 42°C) a second wave of UPR is initiated to handle hyperthermic damage. In MEF cells, however, as a result of membrane perturbation-initiated thermogenesis (SERCA pump uncoupling) the higher local ER temperatures launch a more robust HSR with a downregulated UPR. Because of the higher internal temperature, the second UPR wave starts a few degrees lower in MEF cells, but in both cell lines this second phase of stress response is accompanied by mitigation of HSR.

Previously, we reported the molecular details of cellular “eustress”, wherein CHO cells adapted to mild heat by maintaining membrane homeostasis, activating lipid remodeling, and redistributing chaperone proteins (Peksel et al., 2017). Recent studies have revealed an alternative ER stress activation known as lipid bilayer stress (LBS) which elicits an UPR^LBS^ independently from the classical UPR^PT^ which is responsible for managing proteotoxicity and restoring protein homeostasis. Thibault et al. proposed that LBS serves as a gateway between acute and chronic ER stress (Shyu et al., 2019). Our results suggest that there could be two waves of UPR. UPR^LBS^ is switched on to maintain membrane integrity in U2OS cells as a result of mild heat-induced ER membrane perturbation, while a greater temperature increase would initiate UPR^PT^, as was observed in both cell lines above 42°C (**Fig. 1B**). Most of the data on UPR^LBS^ were obtained from treatments resulting in increased membrane stiffness or decreased membrane fluidity via lipotoxicity, palmitic acid supplementation, or inositol depletion (Celik et al., 2023; Xu and Taubert, 2021). However, there are some indications that compounds that increase membrane fluidity like benzyl alcohol, tergitol, and palmitoleic acid, produce an effect similar to heat and can also initiate the UPR^LBS^ independently of unfolded protein accumulation in yeast (Navarro-Tapia et al., 2018). It was suggested that it is crucial to maintain the biophysical properties of the ER membrane, especially its compressibility (Renne and Ernst, 2023). It may be that since heat alters membrane thickness (Kučerka et al., 2011), lateral diffusion (Peksel et al., 2017), bending and stiffness (Pan et al., 2008), and membrane compressibility, UPR^LBS^ is switched on during mild hyperthermia to maintain the biophysical integrity of all cellular membranes including the ER.

The induction of *XBP1* splicing indicates that the IRE1 pathway is activated by mild heat in U2OS cells (**Fig. 1B, Suppl. Fig. 1**). Our super-resolution fluorescence microscopy data showed that mild ER heat stress induced the clustering of IRE1 similarly to the UPR^PT^ observed by Belyy et al. (2022). It was suggested that the pre-assembled, resting state IRE1 dimers form tetramers (dimers of dimers) upon activation, followed by the appearance of massive clusters if IRE1 expression remains high (Belyy et al., 2022). The increased IRE1 cluster size in the current experiments demonstrates activation of the IRE1 pathway upon mild hyperthermia, which was also confirmed by the downstream induction of XBP1 protein.

In vivo experiments and MD simulations showed that the architecture of the IRE1 molecule could induce compression of the lipid bilayer making it capable of sensing the physical properties of the membrane environment which determine the lifetimes of the dimeric and oligomeric configurations of the ER sensor (Halbleib et al., 2017). While previous studies suggested that IRE1 dimerization was induced by increased saturation and thickening of the bilayer (Halbleib et al., 2017; Kono et al., 2017), our super-resolution experiments demonstrated that the clusterization of IRE1 can also happen in a thinner, more fluid membrane, and that the membrane perturbation induced by mild heat plays a role in the initiation of the UPR^LBS^. The induction of UPR genes upon more severe heat treatment indicates a second wave of responses initiating the classical UPR^PT^ function to either restore proteostasis or induce apoptosis (Almanza et al., 2019).

### Mild heat-induced intracellular thermogenesis

Our novel observation that mild, non-protein denaturing heat induced a HSR in a cell-type-specific manner initiated our studies on intracellular thermogenesis. While the role of mitochondria in non-shivering thermogenesis has already been widely accepted (Fedorenko et al., 2012) the high temperature (Terzioglu et al., 2023) of the organelle is still under debate (Macherel et al., 2021). There has been increasing evidence that the sarco/endoplasmic reticulum Ca^2+^-ATPase (SERCA) pump plays a role in non-shivering thermogenesis in the muscle (Bal et al., 2012), as uncoupling of the SERCA pump results in non-productive activity, increased ATP hydrolysis and heat production (Bal and Periasamy, 2020; Block, 1994).

We hypothesized that a mild environmental temperature increase in MEFs could uncouple the SERCA pump, thereby increasing ER temperature to the levels of HSR initiation. In a reconstituted liposome system, it was shown that optimal functioning of the SERCA pump depended on the conformation of the lipid head groups and the deformation of the bilayer (Andersen and Koeppe, 2007; Caffrey and Feigenson, n.d.; Karlovská et al., 2006). By using the temperature sensitive, ER-specific ER thermo yellow probe and high-content fluorescence imaging we detected temperatures in excess of 48°C in the ER of MEF cells (**Fig. 4**) during mild heating, which could induce protein denaturation in the lumen of the ER and initiate UPR^PT^ (Lepock, 2005).

Further supporting this hypothesis, the ER Ca^2+^ levels drop as a result of mild heat treatment (**Fig. 5**) and TG-inhibition of the SERCA pump suppresses ER thermogenesis (**Fig. 4**). There are few studies reporting elevated ER-temperatures. Using ER thermo yellow, Arai et al. (2014) estimated that the SERCA-uncoupling Ca^2+^ ionophore, ionomycin, caused an ER temperature elevation of 1.7 ± 0.4°C, while using a derivative of this probe, ERthermAC, the induction of thermogenesis by isoproterenol or forskolin raised the temperature from 25°C to 42°C in WT-1 brown adipocyte cells (Kriszt et al., 2017). Recently, it was reported that enhanced Ca^2+^ cycling by activation of the SERCA2b–RyR2 pathway stimulated UCP1**-** and mitochondria-independent thermogenesis in beige adipocytes from mice, humans and pigs (Ikeda et al., 2017). These literature results together with our current finding of an ER temperature increase of 11°C support the hypothesis of a greater role of the ER in thermogenesis.

Although lipid bilayer stress is now established as an ER stress inducer parallel to and independent of proteotoxicity, our understanding of the mechanism of this phenomenon and the signaling pathways involved remains limited, and we are only beginning to understand how the lipidome is influenced by the response of the UPR-ER system to stress. We anticipate that applying the tools of proteomics, metabolomics and lipidomics, will lead to new findings and a more comprehensive understanding. Different environmental, psychological or pathophysiological stresses induce membrane perturbations, which have to be corrected (Török et al., 2014), and this process involves increased metabolic activity to replace or reconfigure lipids and proteins, which could also induce hyperthermia (Oka, 2018).

Taken together, the literature, the data presented here and the observation that the activity of mitochondrial respiratory chain enzymes was found to be maximal at or slightly above 50°C (Chrétien et al., 2018), it may be necessary to reevaluate the concept of homogeneous intracellular temperature distribution, and how membrane integrity can be maintained at these high temperatures. Our previous lipidomics results suggested the existence of a rapid compensatory mechanism involving specific lipid rearrangements by which cells could reestablish normal dynamic membrane phase behavior (Tiszlavicz et al., 2022). The extent of the changes in the lipidome, however, does not suggest a homeoviscous adaptation to very high temperatures; therefore, these studies should be extended to the level of the organelles and membrane proteins potentially involved in thermogenesis. We should also consider the possibility of significant local membrane fluidity alterations because of heterogeneous spatial and temporal temperature fluctuations. Lipids with special properties which could counterbalance local membrane perturbations should also be examined.

The current study revealed that mild hyperthermic stress, like a fever, can lead to a disturbance of ER homeostasis and, depending on the cell type, generate a significant intracellular temperature rise via thermogenic pathways. The stress-induced high internal temperatures could modify the cellular response, and the intrinsic cellular properties could determine the outcome of hyperthermic stress. The data presented here demonstrate that the increased ER temperature initiates a cytosolic HSR, suggesting the existence of ER-associated HSR sensor(s). A role for Hsf1 and other components of the HSR in ER quality control was proposed before to regulate protein folding in the ER and vesicular transport (Liu and Chang, 2008). The literature and our previous studies have lent support to a model in which thermal stress is transduced into a signal at the membrane level (Balogh et al., 2013; Vigh et al., 1998). It was also demonstrated that rapid membrane relocalization of HSPs can maintain membrane fluidity and polymorphism (Horváth et al., 2008); therefore, it is likely that the HSR can also protect ER membranes.

The ER contributes to the production and folding of about a third of cellular proteins, and is thus inextricably linked to the maintenance of cellular homeostasis and the fine balance between health and disease. Our findings may have implications in the development of therapeutic strategies to combat disease in which the UPR was shown to play an important role, like cancer (Horvath et al., 2008), neurodegenerative (Hetz and Saxena, 2017) and ocular diseases (Chen et al., 2023), metabolic diseases like obesity and diabetes (Cnop et al., 2012) and Duchenne’s muscular dystrophy (Fajardo et al., 2017). Knowledge of cell-to-cell variation in response to hyperthermia could also be relevant for preventing heat-stress caused by global warming. The variation in the cellular stress response will affect not only the progression of the disease but also the effectiveness of existing therapeutics (e.g. antibiotics), and also directly affects cell physiology and the ability of the organism to survive extreme heat waves (Kunze et al., 2022).

## Materials and methods

### Cell lines and culture conditions

U2OS (human osteosarcoma) cells were cultured in Dulbecco’s modified Eagle’s medium (DMEM) supplemented with 10% heat-inactivated fetal bovine serum (Gibco, Waltham, MA, USA), 4.5 g/L glucose, 1% L-glutamine, penicillin (200 units/mL) and streptomycin (200 μg/mL) (Sigma-Aldrich, Burlington, MA, USA) at 37°C in a humidified atmosphere of 5% CO_2_. These growing conditions were used for all type of U2OS cells. Cells were routinely tested for mycoplasma contamination using qPCR (MycoQuant Mycoplasma quantification kit, catalog no. MQ-50 AVIDIN, Szeged, Hungary). The U2OS mEOS cell line was created by Belyy et al., and expresses transmembrane IRE1 protein tagged with a C-terminal mEos4b in U2OS IRE1α_KO_ cells. The cell line was a kind gift from Dr. Belyy and its construction is described in Belyy et al. (2020).

To construct the U2OS cell line expressing fluorescently-labeled XBP1 (mNeonGreen), 200 ng linearized pLHCX-XBP1 mNeonGreen plasmid NLS (Addgene plasmid #115971; http://n2t.net/addgene:115971; RRID:Addgene_115971), a gift from David Andrews (Nougarède et al., 2018), was transfected into 10^6^ U2OS cells using the Amaxa Nucleofector System with the Cell Line Nucleofector Kit V (catalog no: VCA-1003, Lonza, Cologne, Germany) utilizing program X-001 according to the manufacturer’s protocol. Twenty-four hours after transfection, the cells were plated at 20% confluence in complete DMEM supplemented with 150 μg/mL hygromycin B (Invitrogen, Thermo Fisher Scientific, Carlsbad CA, US). During selection, the cells were fed with fresh selection medium and resistant colonies arose from single cells were transferred to 96-well plates (Thermo Fisher Scientific, Waltham, MA, USA). Expanded colonies were tested by inducing ER stress with 200 nM thapsigargin for 6 h (Sigma-Aldrich, Burlington, MA, USA) using confocal microscopy (Zeiss LSM800 controlled by the ZEN 2.3 software, Carl Zeiss Microscopy GmbH, Jena, Germany). Colonies without a fluorescent signal in the absence of ER stress, but showing a bright green signal upon induction were considered positive as U2OS XBP1 (UXB3) and used in further experiments.

MEF (mouse embryonic fibroblast) cells were routinely grown in complete Dulbecco’s modified Eagle’s medium (DMEM) supplemented with 10% FBS (Gibco, Waltham, MA, USA), 4.5 g/L glucose, 1% L-glutamine, penicillin (200 units/mL) and streptomycin (200 μg/mL) (Sigma-Aldrich, Burlington, MA, USA) in a 37°C incubator (Galaxy 170S – Eppendorf/New Brunswick) containing 5% CO_2_.

All types of cells were plated at a density of 250 cells/mm^2^ and cultivated in tissue culture plates (VWR, Radnor, PA, USA) or glass-bottom dishes (Greiner Bio-One, Kremsmünster, Austria) 24 hours prior to experiments. Cells were at 80-90% confluence at the time of the experiments, ensuring logarithmic growth during subsequent treatments.

### Fluorescent labeling for measuring ER temperature and Ca^2+^ levels

### Temperature determination with ER thermo yellow (λex/λem 510/591 nm)

U2OS and MEF cells in culture medium were incubated with 250 nM ER thermo yellow (diluted from 1 mM stock solution in DMSO) at 37°C in a humidified atmosphere of 5% CO_2_ for 30 min. After incubation, the cells were washed once with pre-warmed PBS (phosphate buffered saline). ER thermo yellow indicator was a kind gift from Satoshi Arai (National University of Singapore) (Arai et al., 2014). Thapsigargin (Invitrogen, Life Technologies, USA) was used as a positive control and the cells were pre-treated with 1 μM thapsigargin for 1 hour, washed in culture medium, then labeled with ER thermo yellow.

#### Nuclear staining with Hoechst 33342 (λex/λem 350/461 nm)

To facilitate cell segmentation on microscopic images, nuclei were labeled with 1 μg/mL Hoechst 33342 (bisbenzimide trihydrochloride, Sigma-Aldrich, Burlington, MA, USA).

#### Determination of ER Ca^2+^ levels with Mag-Fluo-4 ER (λex/λem 493/517 nm)

Mag-Fluo-4 (M14206, Thermo Fisher Scientific, Waltham, MA, USA) is a low-affinity fluorescent indicator suitable for evaluating the Ca^2+^ concentration in cellular compartments with elevated Ca^2+^ levels, such as the ER (Lebeau et al., 2021). U2OS and MEF cells were incubated in culture medium containing 1 μM Mag-Fluo-4 (1.5 μL of a 1 mM stock solution in DMSO added to 1500 μL of pre-warmed culture medium) at 37°C for 30 minutes in an incubator containing 5% CO_2_. After incubation, cells were washed with pre-warmed PBS, detached with trypsin and resuspended in 500 μL of culture medium.

### Cell fixation

Cells were washed three times with ice-cold PBS, then fixed with 4% PFA (paraformaldehyde) solution for 7 minutes at room temperature, and washed again with ice-cold PBS. Fresh, pre-warmed RPMI 1640 (Sigma Aldrich, Burlington, MA, USA) was added to the cells. RPMI 1640 contains no FBS (Gibco, Waltham, MA, USA) and no phenol-red indicator so culture medium does not interfere with fluorescent imaging.

### Treatments

#### Heat shock experiments

Twenty-four hours prior to heat shock treatment, cells were seeded in tissue culture plates (VWR, Radnor, PA, USA) or glass-bottom dishes (Greiner Bio-One, Kremsmünster, Austria) at a density of 250 cells/mm^2^ in the appropriate complete culture medium. Dishes were sealed with Parafilm and placed into a thermostatically regulated water bath (WNB 7, Memmert) set to different temperatures (39°C, 40°C, 41°C, 42°C, 43°C, 44°C). Samples were analyzed either immediately after the heat shock or after six hours of recovery time at 37°C.

#### Treatment with ER stressors

The culture medium on cells was replaced with medium containing 5 μg/mL tunicamycin (Sigma Aldrich, Burlington, MA, USA) or 100 nM thapsigargin (Invitrogen, Life Technologies, USA), and, after one hour at 37°C, rinsed and replaced with fresh medium.

### RNA isolation and real-time quantitative polymerase chain reaction (RT-qPCR)

U2OS and MEF cells were heat-treated at different temperatures (37°C, 39°C, 40°C, 41°C, 42°C, 43°C, 44°C) or treated with 5 μg/mL tunicamycin or 100 nM thapsigargin for one hour. After one hour of recovery, RNA was isolated utilizing an RNA and protein purification kit (Macherey-Nagel, Düren, Germany) as per the manufacturer’s guidelines. RNA was converted into cDNA using the high-capacity cDNA reverse transcription kit (Thermo Fisher Scientific, Waltham, MA, USA). Each reaction mix consisted of 15 µL mRNA (1 µg), 3 µL primer, 1.5 µL reverse transcriptase, 1.2 µL dNTPs, 3 µL buffer, and 6.3 µL RNase-free water. The reverse transcription process was carried out using a T100 thermocycler (BioRad, Hercules, CA, USA) according to the following protocol: 10 min incubation at 25°C, reverse transcription for 2 h at 37°C, and 5 min of inactivation at 85°C. The resulting cDNA product was diluted 1:20 for use as a template in a 20 μl RT-qPCR reaction containing 9 µL of cDNA, 1 µL of primer mix (250 nM for both forward and reverse primers), and 10 µL of 2x Power SYBR-Green PCR master mix (Thermo Fisher Scientific, Waltham, MA, USA). The reactions were run on a RotorGene 3000 instrument (Qiagen, Hilden, Germany) with the following settings: initial denaturation at 95°C for 10 min, followed by 40 cycles of denaturation at 95°C for 15 seconds, and annealing at 60°C for 60 seconds. Melting curve analysis spanning 50–95°C was performed to confirm the specificity of amplification. Primer sequences utilized in RT-qPCR reactions are listed in Supplementary Table 3. Mouse *GAPDH* was used for MEF cell samples while human *GAPDH* (glyceraldehyde 3-phosphate dehydrogenase), and human *RPL27* (ribosomal protein L27) genes were used for the U2OS cell line as an internal normalization control. Relative gene expression levels were determined using the ΔΔCt method.

### Flow cytometry analysis

#### Measuring XBP1 protein levels

U2OS cells stably expressing fluorescently tagged XBP1 (mNeonGreen) protein were seeded in 35 mm glass-bottom dishes (Greiner Bio-One, Kremsmünster, Austria) at a density of 250 cells/mm^2^ 48 hours prior to experiments. Cells were heat-treated at 40°C and 42°C, or treated with tunicamycin (5 μg/mL) for 1 h, followed by 6 h of recovery at 37°C, then trypsinized and washed with culture medium. All cell analyses were performed using a BD Accuri C6 flow cytometer (BD Biosciences, CA, USA) equipped with standard laser and filter setup. Data were obtained with linear amplification of the forward (FSC) and side scatter (SSC) light signals. Fluorescence signals emitted by the mNeonGreen-tagged XBP1 proteins were measured by FL1 detector for green fluorescence with logarithmic amplification. The light emission from 10,000 cells per sample was analyzed, and the frequency distribution histograms were created with the web-based program, floreada.io (https://floreada.io) in FL1. The fluorescence intensity values of the treated cells were compared to the intensity values of the untreated cells (maintained at 37°C).

#### Measuring ER Ca^2+^ levels

Mag-fluo-4 labeled MEF and U2OS cells were seeded in 35 mm dishes (Greiner Bio-One, Kremsmünster, Austria) at a density of 250 cells/mm^2^ 24 hours prior to experiments. Cells were trypsinized and washed with culture media. Fluorescence intensity of Mag-fluo-4-loaded cells was measured using a BD Accuri C6 flow cytometer (BD Biosciences, CA, USA) with an FL1 detector for green fluorescence and logarithmic amplification. The light emission from 10,000 cells per sample was analyzed, and the frequency distribution histograms were created with the web-based program, floreada.io (https://floreada.io) in FL1. Cells were heated to 40°C in 10 minutes and the mean fluorescence intensity of the treated cells was compared to the values of the untreated cells (maintained at 37°C). Cells were also pre-treated with the SERCA pump inhibitor, 1 μM thapsigargin, for one hour.

### Fluorescence microscopy assays

#### ER temperature measurements by ER thermo yellow fluorescence

Fluorescence intensity changes of ER thermo yellow-labeled cells were monitored using a Cell-Discoverer 7 fluorescence microscope (Zeiss, Jena, Germany) equipped with the Axiocam 506 imaging device, for controlling light, temperature and CO_2_. For the ER thermo yellow indicator we used λ_ex_528/λ_em_553 and for the Hoechst stain, λ_ex_348/λ_em_ 455, with 25% and 6% light source intensity, respectively. One image was acquired at 37°C and within 10 min, another image at 40°C to reduce the possibility of photobleaching. Approximately 1000 cells were acquired from one image, and they were accurately segmented for further analysis utilizing Matlab (MathWorks Inc., Massachusetts, US), ImageJ (Rasband, W.S., ImageJ, U. S. National Institutes of Health, Bethesda, Maryland, USA), and CellProfiler (Broad Institute, Cambridge, MA, USA). Fluorescence intensity values of individual cells were averaged, and the average intensity of the heat-treated sample (F) was normalized to the intensity value of the pre-heated sample (F0), resulting in the F/F0 ratio in which F/F0 = 1 meant no change in ER temperature, F/F0 < 1 meant increased temperature, and F/F0 > 1 meant decreased temperature). Changes in F/F0 of fixed cells gives information about the external temperature (37-40°C) change, and any additional change in live cells provides information about the changes of the ER temperature. Increasing ambient temperature from 37°C to 40°C resulted in a 10% decrease in the F/F0 ratio of fixed cells, which was used to calibrate the temperature changes in live cells (Fig. 4C).

#### Super-resolution microscopy

Super-resolution fluorescence measurements were performed on a custom-built inverted microscope system based on a Nikon Eclipse Ti-E frame. Images were captured under epi-illumination with an oil immersion objective (Nikon CFI Apo 100x, NA = 1.49). Fluorescent proteins were photoconverted by a 405 nm laser (Nichia, 60 mW) and were localized by a readout laser operating at 561 nm (Cobolt Jive, 300 mW). A fluorescent filter set (Semrock, LF405/488/561/635-A-000) and an additional emission filter (Semrock, FF01-600/37) were used to separate the excitation and emission wavelengths. Images were captured by an EMCCD digital camera (Andor iXon3 897, 512x512 pixels, 16 µm pixel size). Typically, 3,000-6,000 frames were captured with an exposure time of 100 ms (until no more photoconversion was detectable). Image stacks were evaluated with the rainSTORM localization software (Rees et al., 2013). Only localizations with precisions of <45 nm were used for further analysis.

Density-based spatial clustering of applications with density-based cluster analysis (DBSCAN) was performed on the localization datasets for quantitative evaluation. After thorough tests (Suppl. Fig. 4), the input parameters for this algorithm were set to ε = 45 (the maximum distance between two adjacent points) and N = 10 (the minimum number of points that form a cluster). Cluster parameters like cluster area and number of localizations per cluster were calculated and saved for further analysis.

## Authors contribution

Conceptualization: B.D., I.G., Z.T.; Investigation: B.D., I.G., Z.R., Á.H., P.B.; Methodology: M.E.T., Á.C., P.B., Á.T., G.T.; Validation: B.D., I.G., Z.T., M.E.; Formal analysis: B.D., I.G., M.P., G.B., P.B.; Writing – original draft preparation: B.D., I.G., Z.T.; Writing – review & editing: M.P., G.B., M.E.T., M.E., L.V.; Visualisation: I.G., M.P., P.B.; Funding acquisition: Z.T.

