## Supplement Material for "Mild hyperthermia-induced thermogenesis in the endoplasmic reticulum defines stress response mechanisms"

| **MEF** | 37°C | 39°C | 40°C | 41°C | 42°C | 43°C | 44°C |
|---|---|---|---|---|---|---|---|
| *Hsp25* | 100 | 215.348 | 3530.17 | 30443.7 | 44882.2 | 5623.25 | 6370.49 |
| *Hsp60* | 100 | 116.473 | 151.747 | 178.18 | 148.11 | 86.8541 | 96.1483 |
| *Hsp70* | 100 | 987.194 | 17026.8 | 64359.1 | 94445.2 | 30163.6 | 33546.1 |
| *Hsp90aa1* | 100 | 143.396 | 192.964 | 288.786 | 265.737 | 97.716 | 95.705 |
| *Hsp90ab1* | 100 | 110.957 | 129.984 | 119.748 | 116.205 | 108.423 | 108.925 |
| *Hsf1* | 100 | 84.6745 | 92.7659 | 76.8438 | 82.169 | 91.5945 | 79.1869 |
| *Ire1a* | 100 | 106.93 | 102.693 | 101.396 | 104.247 | 105.214 | 103.526 |
| *Ire1b* | 100 | 102.811 | 114.472 | 104.729 | 104.729 | 110.191 | 105.458 |
| *Xbp1_total* | 100 | 131.039 | 127.603 | 94.8246 | 63.2878 | 99.7692 | 99.0801 |
| *Xbp1_us* | 100 | 120.025 | 145.229 | 109.682 | 51.0506 | 30.2848 | 31.9377 |
| *Xbp1_s* | 100 | 67.8694 | 44.7254 | 29.6787 | 45.5072 | 135.426 | 138.591 |
| *Bip* | 100 | 124.258 | 127.899 | 104.488 | 81.0378 | 93.088 | 93.7354 |
| *Atf6* | 100 | 93.9523 | 116.339 | 89.0899 | 84.0896 | 123.114 | 130.435 |
| *Perk* | 100 | 101.63 | 106.807 | 97.716 | 97.716 | 105.214 | 105.458 |
| *Chop* | 100 | 102.337 | 91.4888 | 67.6737 | 87.0551 | 64.3197 | 53.8369 |
| *Grp94* | 100 | 106.437 | 119.886 | 100.463 | 96.1483 | 99.3092 | 106.437 |
| *Atf4* | 100 | 43.4271 | 39.3199 | 28.5851 | 23.0579 | 99.3092 | 92.445 |

| **U2OS** | 37°C | 39°C | 40°C | 41°C | 42°C | 43°C | 44°C |
|---|---|---|---|---|---|---|---|
| HSP27 | 100 | 113.158 | 143.23 | 199.538 | 246.514 | 129.235 | 140.932 |
| HSP60 | 100 | 102.456 | 115.535 | 155.473 | 233.216 | 121.982 | 165.29 |
| HSP70 | 100 | 295.878 | 703.719 | 1163.18 | 2013.55 | 1758.98 | 2138.21 |
| HSP90aa1 | 100 | 99.8845 | 119.61 | 141.421 | 185.104 | 107.923 | 119.61 |
| HSP90ab1 | 100 | 113.945 | 125.266 | 142.076 | 147.597 | 118.646 | 133.947 |
| HSF1 | 100 | 179.212 | 203.967 | 221.914 | 180.042 | 133.793 | 153.865 |
| IRE1a | 100 | 74.4839 | 87.1557 | 119.196 | 128.194 | 72.6986 | 85.362 |
| XBP1_total | 100 | 148.624 | 164.148 | 173.708 | 153.51 | 87.0551 | 101.983 |
| XBP1_us | 100 | 138.191 | 141.748 | 138.191 | 117.283 | 9.06636 | 26.4255 |
| XBP1_s | 100 | 201.391 | 293.494 | 262.685 | 334.808 | 1024.37 | 873.415 |
| BIP | 100 | 92.7659 | 99.1946 | 99.539 | 95.5945 | 93.088 | 94.2785 |
| ATF6 | 100 | 85.362 | 101.748 | 86.0551 | 95.1538 | 90.5425 | 93.6272 |
| PERK | 100 | 128.491 | 147.257 | 144.727 | 138.351 | 94.3874 | 115.003 |
| CHOP | 100 | 372.782 | 406.992 | 349.026 | 279.917 | 69.7372 | 104.85 |
| ATF4 | 100 | 92.445 | 99.539 | 89.9171 | 88.0666 | 102.337 | 104.972 |

**Supplementary Table 1.** Transcription profile of important HSP and UPR genes in MEF and U2OS cells in response to heat treatment at different temperatures (37°C, 39°C, 40°C, 41°C, 42°C, 43°C, 44°C) for one hour. Expression of target genes in treated cells was compared to non-treated control cells kept at optimal growing temperature of 37°C (results are given as a percentage, where non-treated cells = 100%). Student's t-test was used for statistical comparisons ($p$<0.05; significant change indicated with blue).

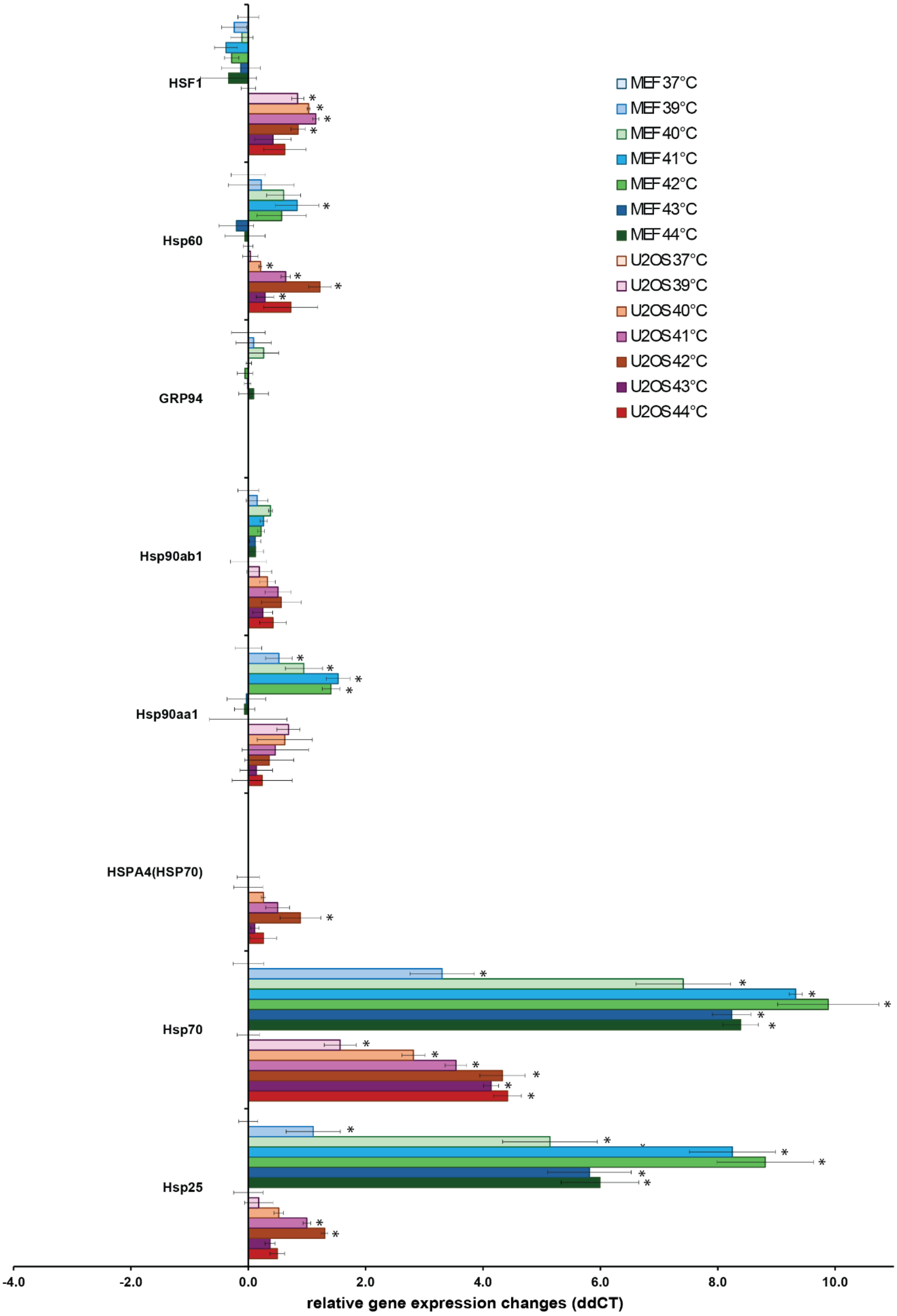
HSF1
Hsp60
GRP94
Hsp90ab1
Hsp90aa1
HSPA4(HSP70)
Hsp70
Hsp25
MEF 37°C
MEF 39°C
MEF 40°C
MEF 41°C
MEF 42°C
MEF 43°C
MEF 44°C
U2OS 37°C
U2OS 39°C
U2OS 40°C
U2OS 41°C
U2OS 42°C
U2OS 43°C
U2OS 44°C
-4.0
-2.0
0.0
2.0
4.0
6.0
8.0
10.0
relative gene expression changes (ddCT)

**Supplementary Figure 1. Relative heat-shock protein (HSP) gene expression changes (ddCT) in MEF and U2OS cells in response to heat treatment at different temperatures (39°C, 40°C, 41°C, 42°C, 43°C, 44°C) for one hour.** The relative expression of different HSP genes was studied using RT-qPCR ($n$ = 3 biological repetitions). The expression of target genes in treated cells was compared to non-treated control cells kept at the optimal growing temperature of 37°C. Student's t-test was used for statistical comparisons (*$p$<0.05).

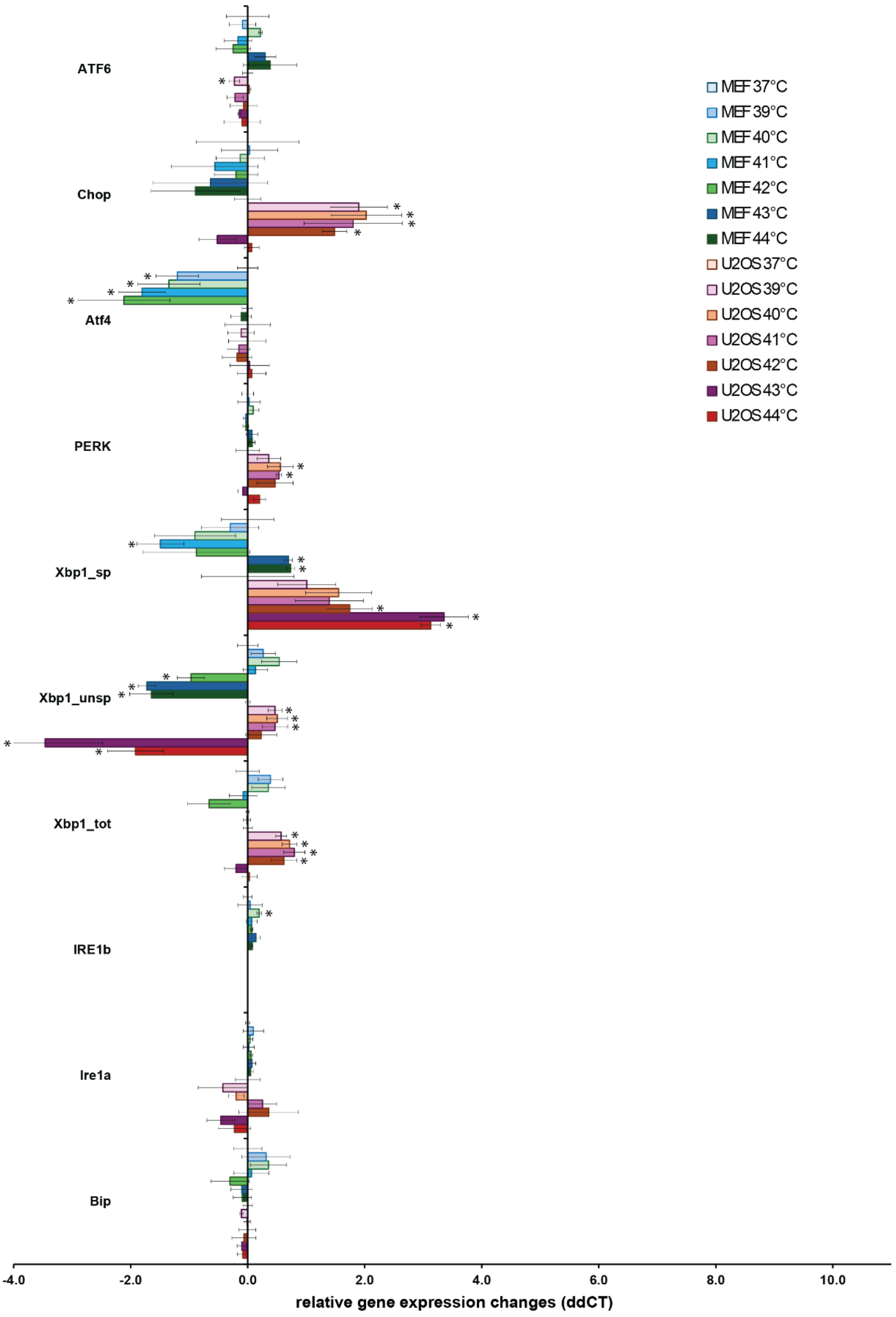

ATF6
Chop
Atf4
PERK
Xbp1_sp
Xbp1_unsp
Xbp1_tot
IRE1b
Ire1a
Bip
MEF 37°C
MEF 39°C
MEF 40°C
MEF 41°C
MEF 42°C
MEF 43°C
MEF 44°C
U2OS 37°C
U2OS 39°C
U2OS 40°C
U2OS 41°C
U2OS 42°C
U2OS 43°C
U2OS 44°C
-4.0
-2.0
0.0
2.0
4.0
6.0
8.0
10.0
relative gene expression changes (ddCT)

**Supplementary Figure 2. Relative unfolded protein response (UPR) gene expression changes (ddCT) in MEF and U2OS cells in response to heat treatment at different temperatures (39°C, 40°C, 41°C, 42°C, 43°C, 44°C) for one hour.** The relative expression of different UPR genes was studied using RT-qPCR (*n* = 3 biological repetitions). The expression of target genes in treated cells was compared to non-treated control cells kept at the optimal growing temperature of 37°C. Student's t-test was used for statistical comparisons (**p*<0.05).

MEF

| | Hsp25 | Hsp70 | Hsp90aa1 | Hsp90ab1 | Hsp60 | HSF1 | Bip | Ire1a | Xbp1_tot | Xbp1_us | Xbp1_sp | PERK | Atf4 | Chop | ATF6 |
|---|---|---|---|---|---|---|---|---|---|---|---|---|---|---|---|
| Hsp25 | 1.000 | 0.961 | 0.597 | 0.466 | 0.395 | -0.274 | -0.309 | -0.006 | -0.523 | -0.360 | -0.305 | -0.048 | -0.476 | -0.326 | -0.085 |
| Hsp70 | 0.961 | 1.000 | 0.490 | 0.483 | 0.337 | -0.297 | -0.247 | 0.044 | -0.409 | -0.409 | -0.229 | 0.006 | -0.418 | -0.342 | 0.049 |
| Hsp90aa1 | 0.597 | 0.490 | 1.000 | 0.636 | 0.802 | -0.123 | -0.045 | -0.045 | -0.354 | 0.406 | -0.823 | -0.201 | -0.906 | -0.067 | -0.465 |
| Hsp90ab1 | 0.466 | 0.483 | 0.636 | 1.000 | 0.752 | 0.241 | 0.318 | -0.154 | 0.079 | 0.307 | -0.387 | 0.311 | -0.496 | -0.157 | 0.111 |
| Hsp60 | 0.395 | 0.337 | 0.802 | 0.752 | 1.000 | 0.276 | 0.019 | -0.348 | -0.265 | 0.439 | -0.727 | -0.011 | -0.735 | -0.159 | -0.209 |
| HSF1 | -0.274 | -0.297 | -0.123 | 0.241 | 0.276 | 1.000 | -0.107 | -0.185 | 0.064 | 0.102 | 0.091 | 0.251 | 0.147 | -0.278 | 0.193 |
| Bip | -0.309 | -0.247 | -0.045 | 0.318 | 0.019 | -0.107 | 1.000 | -0.081 | 0.766 | 0.603 | 0.047 | 0.350 | 0.179 | 0.513 | -0.013 |
| Ire1a | -0.006 | 0.044 | -0.045 | -0.154 | -0.348 | -0.185 | -0.081 | 1.000 | 0.126 | -0.138 | 0.166 | 0.059 | -0.019 | 0.049 | -0.054 |
| Xbp1_tot | -0.523 | -0.409 | -0.354 | 0.079 | -0.265 | 0.064 | 0.766 | 0.126 | 1.000 | 0.451 | 0.285 | 0.322 | 0.387 | 0.319 | 0.245 |
| Xbp1_us | -0.360 | -0.409 | 0.406 | 0.307 | 0.439 | 0.102 | 0.603 | -0.138 | 0.451 | 1.000 | -0.599 | -0.113 | -0.366 | 0.421 | -0.399 |
| Xbp1_sp | -0.305 | -0.229 | -0.823 | -0.387 | -0.727 | 0.091 | 0.047 | 0.166 | 0.285 | -0.599 | 1.000 | 0.517 | 0.830 | -0.040 | 0.503 |
| PERK | -0.048 | 0.006 | -0.201 | 0.311 | -0.011 | 0.251 | 0.350 | 0.059 | 0.322 | -0.113 | 0.517 | 1.000 | 0.343 | -0.143 | 0.267 |
| Atf4 | -0.476 | -0.418 | -0.906 | -0.496 | -0.735 | 0.147 | 0.179 | -0.019 | 0.387 | -0.366 | 0.830 | 0.343 | 1.000 | 0.030 | 0.466 |
| Chop | -0.326 | -0.342 | -0.067 | -0.157 | -0.159 | -0.278 | 0.513 | 0.049 | 0.319 | 0.421 | -0.040 | -0.143 | 0.030 | 1.000 | -0.253 |
| ATF6 | -0.085 | 0.049 | -0.465 | 0.111 | -0.209 | 0.193 | -0.013 | -0.054 | 0.245 | -0.399 | 0.503 | 0.267 | 0.466 | -0.253 | 1.000 |

U2OS

| | Hsp25 | Hsp70 | Hsp90aa1 | Hsp90ab1 | Hsp60 | HSF1 | Bip | Ire1a | Xbp1_tot | Xbp1_unsp | Xbp1_sp | PERK | Atf4 | Chop | ATF6 |
|---|---|---|---|---|---|---|---|---|---|---|---|---|---|---|---|
| Hsp25 | 1.000 | 0.650 | 0.096 | 0.537 | 0.816 | 0.590 | 0.009 | 0.464 | 0.505 | 0.200 | 0.158 | 0.545 | -0.369 | 0.402 | -0.040 |
| Hsp70 | 0.650 | 1.000 | 0.066 | 0.643 | 0.723 | 0.451 | -0.101 | 0.156 | 0.058 | -0.419 | 0.790 | 0.197 | 0.063 | -0.052 | -0.053 |
| Hsp90aa1 | 0.096 | 0.066 | 1.000 | 0.402 | -0.014 | 0.425 | 0.450 | 0.111 | 0.536 | 0.321 | -0.024 | 0.260 | 0.397 | 0.287 | 0.188 |
| Hsp90ab1 | 0.537 | 0.643 | 0.402 | 1.000 | 0.718 | 0.609 | 0.525 | 0.668 | 0.431 | 0.151 | 0.306 | 0.481 | 0.401 | 0.137 | 0.351 |
| Hsp60 | 0.816 | 0.723 | -0.014 | 0.718 | 1.000 | 0.450 | 0.049 | 0.552 | 0.215 | 0.044 | 0.298 | 0.378 | -0.087 | 0.091 | 0.135 |
| HSF1 | 0.590 | 0.451 | 0.425 | 0.609 | 0.450 | 1.000 | 0.107 | 0.287 | 0.724 | 0.468 | 0.052 | 0.743 | -0.103 | 0.719 | -0.009 |
| Bip | 0.009 | -0.101 | 0.450 | 0.525 | 0.049 | 0.107 | 1.000 | 0.547 | 0.362 | 0.298 | -0.201 | 0.254 | 0.554 | -0.036 | 0.632 |
| Ire1a | 0.464 | 0.156 | 0.111 | 0.668 | 0.552 | 0.287 | 0.547 | 1.000 | 0.397 | 0.465 | -0.306 | 0.452 | 0.169 | 0.145 | 0.355 |
| Xbp1_tot | 0.505 | 0.058 | 0.536 | 0.431 | 0.215 | 0.724 | 0.362 | 0.397 | 1.000 | 0.749 | -0.294 | 0.805 | -0.094 | 0.841 | 0.052 |
| Xbp1_unsp | 0.200 | -0.419 | 0.321 | 0.151 | 0.044 | 0.468 | 0.298 | 0.465 | 0.749 | 1.000 | -0.759 | 0.598 | -0.142 | 0.758 | 0.153 |
| Xbp1_sp | 0.158 | 0.790 | -0.024 | 0.306 | 0.298 | 0.052 | -0.201 | -0.306 | -0.294 | -0.759 | 1.000 | -0.219 | 0.257 | -0.323 | -0.101 |
| PERK | 0.545 | 0.197 | 0.260 | 0.481 | 0.378 | 0.743 | 0.254 | 0.452 | 0.805 | 0.598 | -0.219 | 1.000 | -0.295 | 0.753 | 0.086 |
| Atf4 | -0.369 | 0.063 | 0.397 | 0.401 | -0.087 | -0.103 | 0.554 | 0.169 | -0.094 | -0.142 | 0.257 | -0.295 | 1.000 | -0.356 | 0.527 |
| Chop | 0.402 | -0.052 | 0.287 | 0.137 | 0.091 | 0.719 | -0.036 | 0.145 | 0.841 | 0.758 | -0.323 | 0.753 | -0.356 | 1.000 | -0.137 |
| ATF6 | -0.040 | -0.053 | 0.188 | 0.351 | 0.135 | -0.009 | 0.632 | 0.355 | 0.052 | 0.153 | -0.101 | 0.086 | 0.527 | -0.137 | 1.000 |

**Supplementary Table 2. Heat map representation of correlation data between the heat-shock protein (HSP) and unfolded protein response (UPR) genes of MEF and U2OS cells following exposure to heat.** Pearson correlation analysis was conducted to examine the relationship between the induced HSP and UPR genes. Color code: -1 = total negative linear correlation (green); 0 = no correlation (yellow); +1 = total positive correlation (red).

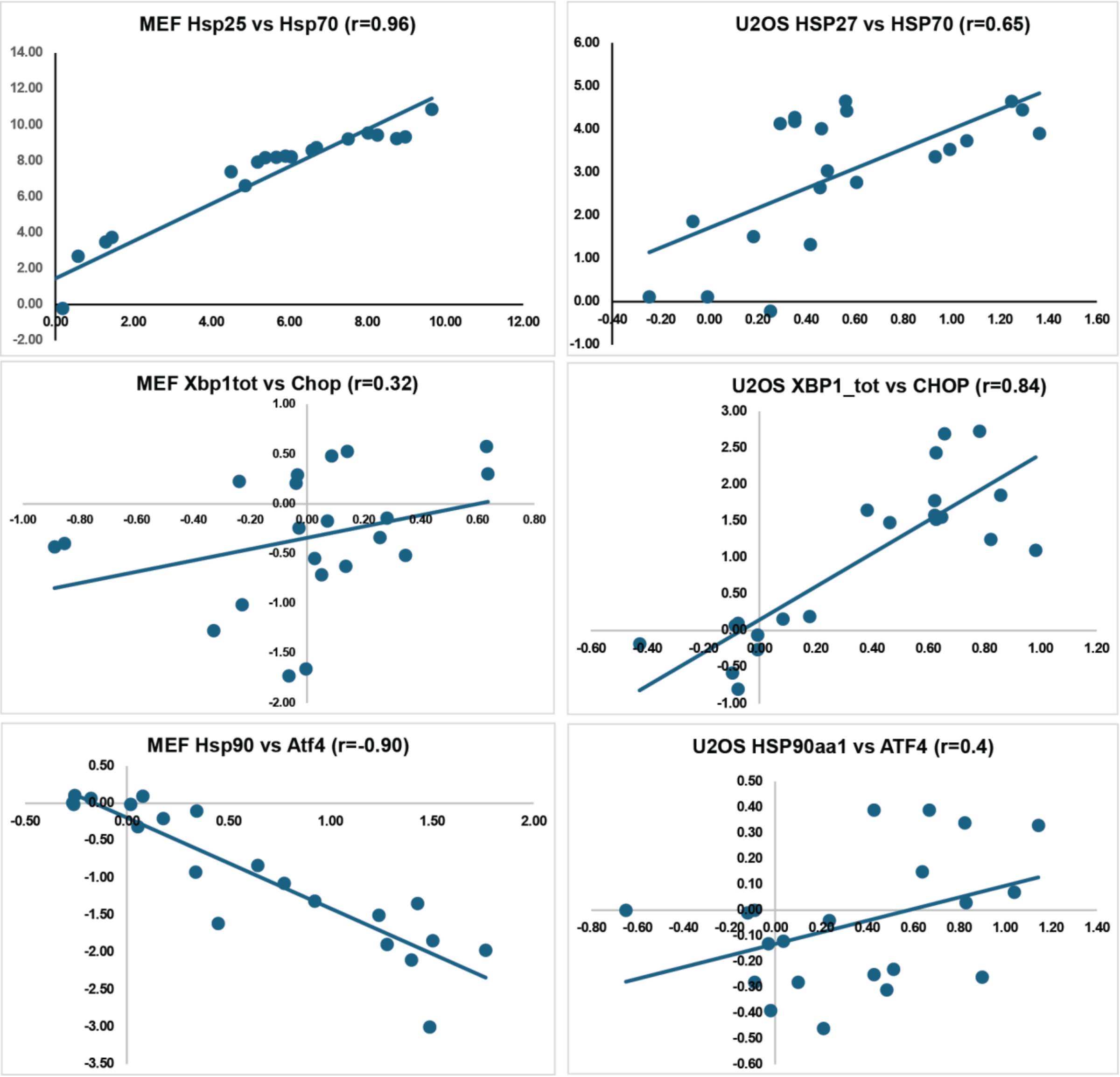


**Supplementary Figure 3. Correlation between HSR and UPR genes in MEF and U2OS cells.** Circles in scatter plots represent individual samples; n = 3 (independent biological replicates) at each applied temperatures from 37 to 44°C. Solid lines show linear regressions, r values represent Pearson correlation coefficients.

| Mouse gene | | forward primer | reverse primer |
|---|---|---|---|
| **1** | ***Gapdh*** | GGGTTCCTATAAATACGGACTGC | CCATTTTGTCTACGGGACGA |
| **2** | ***Xbp1_total*** | TGGCCGGGTCTGCTGAGTCCG | GTCCATGGGAAGATGTTCTGG |
| **3** | ***Xbp1_us*** | CAGCACTCAGACTATGTGCA | GTCCATGGGAAGATGTTCTGG |
| **4** | ***Xbp1_s*** | CTGAGTCCGAATCAGGTGCAG | GTCCATGGGAAGATGTTCTGG |
| **5** | ***Bip*** | TTCAGCCAATTATCAGCAAACTCT | TTTTCTGATGTATCCTCTTCACCAGT |
| **6** | ***Ire1a/Ern1*** | CTGGCTTCTCATAGGACACCAT | TCTCGATGTTTGGGCAGGTT |
| **7** | ***Ire1b/Ern2*** | CCATGAGGAACAAGAAGCACC | AGGAAGTTGGCCTAGTGTCTG |
| **8** | ***Perk*** | TCCTGCTTTGCATCGTAGCC | CAGACTCCTTCCGCTGCCTG |
| **9** | ***Atf4*** | CTAAGCCATGGCGCTCTTCA | GCTGGATTCGAGGAATGTGC |
| **10** | ***Atf6*** | CCAGATGAAGACTGGGAGTCG | GCTGCATCAAAGTGCACATCA |
| **11** | ***Chop*** | CCACCACACCTGAAAGCAGAA | AGGTGAAAGGCAGGGACTCA |
| **12** | ***Hsf1*** | GGGAAACAGGAGTGTATGGACT | CTTGTTGACAACTTTTTGCTGCT |
| **13** | ***Hsp25*** | ATCCCCTGAGGGCACACTTA | GGAATGGTGATCTCCGCTGAC |
| **14** | ***Hsp60*** | CACAGTCCTTCGCCAGATGAG | CTACACCTTGAAGCATTAAGGCT |
| **15** | ***Hsp70*** | GAGATCGACTCTCTGTTCGAGG | GCCCGTTGAAGAAGTCCTG |
| **16** | ***Hsp90aa1*** | GACGCTCTGGATAAAATCCGTT | TGGGAATGAGATTGATGTGCAG |
| **17** | ***Hsp90ab1*** | AAACAAGGAGATTTTCCTCCGC | CCGTCAGGCTCTCATATCGAAT |
| **Human gene** | | **forward primer** | **reverse primer** |
| **18** | ***GAPDH*** | ACAACTTTGGTATCGTGGAAGG | GCCATCACGCCACAGTTTC |
| **19** | ***RPL27*** | CGCAAAGCTGTCATCGTG | GTCACTTTGCGGGGGTAG |
| **20** | ***XBP1_total*** | TGAAAAACAGAGTAGCAGCTCAGA | CCCAAGCGCTGTCTTAACTC |
| **21** | ***XBP1_us*** | CAGACTACGTGCACCTCTGC | CTGGGTCCAAGTTGTCCAGAAT |
| **22** | ***XBP1_s*** | GCTGAGTCCGCAGCAGGT | CTGGGTCCAAGTTGTCCAGAAT |
| **23** | ***BIP*** | GACCACCTACTCCTGCGTC | GTGAAGGCGACATAGGACGG |
| **24** | ***IRE1a/ERN1*** | AATTGTGTACCGGGGCATGT | TCACGGTCTGCGAAGCTAAA |
| **25** | ***IRE1b/ERN2*** | CGCTGAGTCCACAGGTTCATA | GTCTGCTTGCTTAGTGCGTG |
| **26** | ***PERK*** | GAGAGCAGGAGCCTCGG | GATAATTACTAATGACCTGCCGC |
| **27** | ***ATF4*** | TTAAGCCATGGCGCTTCTCA | GCTGGAATCGAGGAATGTGCT |
| **28** | ***ATF6*** | ACCCGTATTCTTCAGGGTGC | TCCCTGAGTTCCTGCTGATAC |
| **29** | ***CHOP*** | ACCTGAGGAGAGAGTGTTCA | TCAGTCTGGAAAAGCACATCT |
| **30** | ***HSF1*** | AGCCTGGCCAGTATCCAAGA | AGCTGCTTCCCTGAATCCG |
| **31** | ***HSP27*** | GTCCCTGGATGTCAACCACT | GACTGGGATGGTGATCTCGT |
| **32** | ***HSP60*** | ACTCGGAGGCGGAAGAAAAA | GTAACCGAAGCATTTCTCGGC |
| **33** | ***HSP70/HSPA1A*** | CCCCACCATTGAGGAGGTAG | AGGAAATGCAAAGTCTTGAAGCTC |
| **34** | ***HSP90AA1*** | GAATACCCGCGCGACCGT | TTAACAGGTGCCCTGCTTCTCAG |
| **35** | ***HSP90AB1*** | GGGTATCGGAAAGCAAGCCT | GCACTTCCTCAGGCATCTTGAA |

**Supplementary Table 3.** Primer sequences utilized in RT-qPCR reactions
(Mouse Xbp1 primers are from Oslowski CM, Urano F. Measuring ER stress and the unfolded protein response using mammalian tissue culture system. Methods Enzymol. 2011;490:71-92. doi: 10.1016/B978-0-12-385114-7.00004-0. PMID: 21266244; PMCID: PMC3701721.)

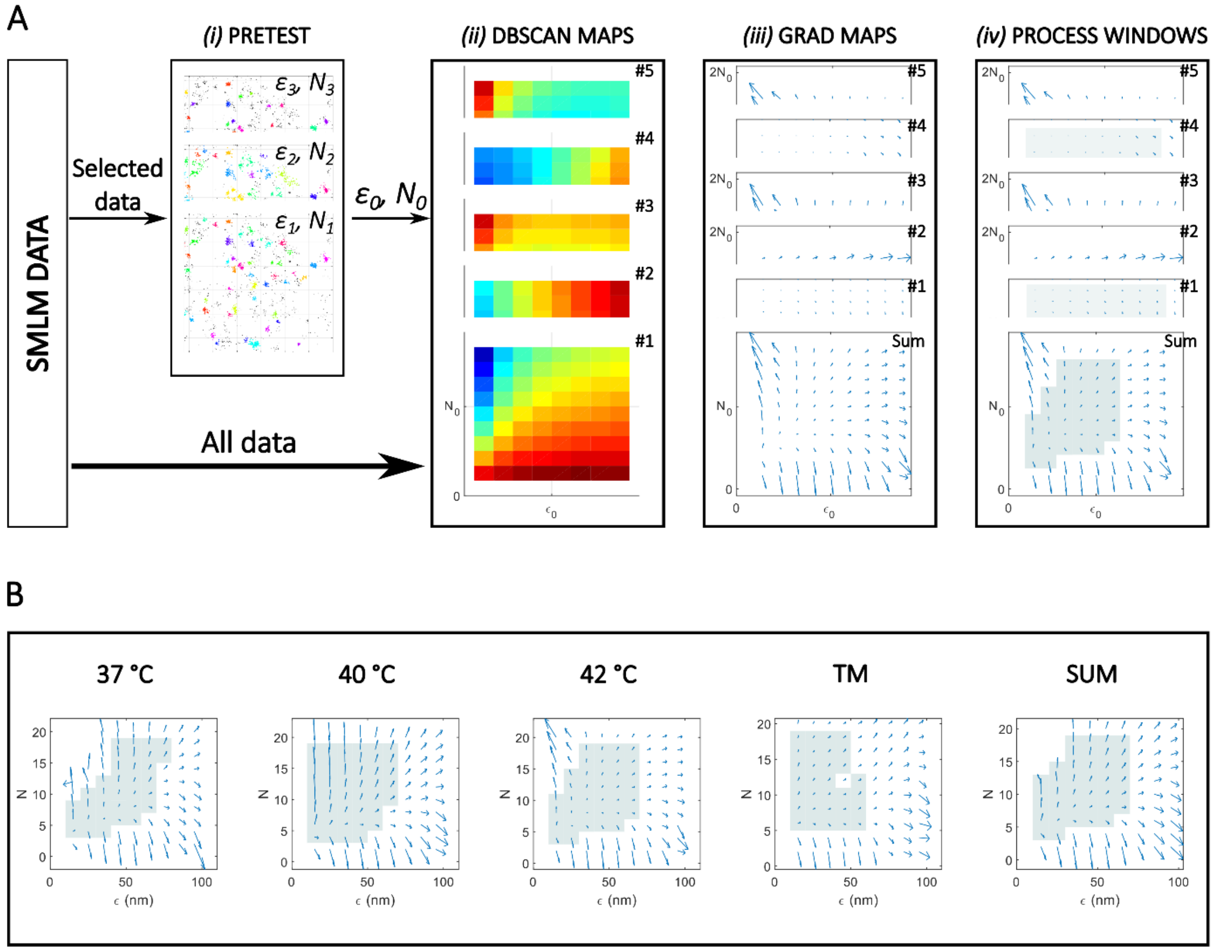


**Supplementary Figure 4. Selection procedure of the optimum DBSCAN parameters.** The DBSCAN cluster analyzer algorithm uses two input parameters, a distance ($\varepsilon$) and a minimum number of points ($N$) values. Ideally a slight change in such parameters do not significantly affect the feature of the final clustered images, therefore quantitative analysis can be associated with real treatments and changes in the sample. Taking into account this condition, the optimum $\varepsilon$ and $N$ values were selected via a four-step process (A). First, a pretest cluster analysis (i) was performed on the same type of samples with several $\varepsilon$ and $N$ values to visually find the optimum clustered images. In the present work $\varepsilon_0$=55 nm and $N_0$=12 were selected. Next, DBSCAN cluster analyses (ii) were performed using the parameter window with size of $2\varepsilon_0$ and $2N_0$, and DBSCAN maps were generated using five merit functions (cluster area mean and standard deviation, localization number per cluster mean and standard deviation and number of clusters). Next, gradient maps (iii) for all DBSCAN maps were generated. At the same time the normalized gradient map of the sum was also calculated. Finally, a process window (iv) was selected on the gradient maps, where the sum gradient value was less than 0.25.

The process was repeated for all the treated, the control and the sum images (B). The optimum $\varepsilon$-$N$ value was defined as the center of such process windows. Based on these results, the originally selected parameter values were slightly modified to $\varepsilon$=45 nm and $N$=10, and all the cluster analyses were performed under such conditions.
